# Postnatal fluoxetine treatment alters perineuronal net formation and maintenance in the hippocampus

**DOI:** 10.1101/2020.10.12.336941

**Authors:** Sourish Mukhopadhyay, Ashmita Chatterjee, Praachi Tiwari, Utkarsha Ghai, Vidita A. Vaidya

## Abstract

Elevation of serotonin via postnatal fluoxetine (PNFlx) treatment during critical temporal windows is hypothesized to perturb the development of limbic circuits thus establishing a substratum for persistent disruption of mood-related behavior. We examined the impact of PNFlx treatment on the formation and maintenance of perineuronal nets (PNNs), extracellular matrix (ECM) structures that deposit primarily around inhibitory interneurons, and mark the closure of critical period plasticity. PNFlx treatment evoked a significant decline in PNN number, with a robust reduction in PNNs deposited around parvalbumin (PV) interneurons, within the CA1 and CA3 hippocampal subfields at postnatal day 21 in Sprague-Dawley rat pups. While the reduction in CA1 subfield PNN number was still observed in adulthood, we observed no change in colocalization of PV-positive interneurons with PNNs in the hippocampi of adult PNFlx animals. PNFlx treatment did not alter hippocampal parvalbumin, calretinin, or reelin-positive neuron numbers in PNFlx animals at P21 or in adulthood. We did observe a small, but significant increase in somatostatin (SST)-positive interneurons in the DG subfield of PNFlx-treated animals in adulthood. This was accompanied by altered GABA-A receptor subunit composition, increased dendritic complexity of apical dendrites of CA1 pyramidal neurons, and enhanced neuronal activation revealed by increased c-Fos-positive cell numbers within hippocampi of PNFlx-treated animals in adulthood. These results indicate that PNFlx treatment alters the developmental trajectory of PNNs within the hippocampus, raising the possibility of a disruption of critical period plasticity and the establishment of an altered excitation-inhibition balance within this key limbic brain region.

**Significance Statement:** Clinical and preclinical studies indicate that developmental exposure to fluoxetine programs persistent dysregulation of mood-related behaviors. This is hypothesized to involve the disruption of the normal development of key brain regions, such as the hippocampus that regulate mood behaviors. We show that postnatal exposure to fluoxetine alters hippocampal perineuronal nets (PNNs), extracellular matrix structures that regulate plasticity and mark the closure of critical periods. The decline in PNNs is noted in early postnatal life, and persists into adulthood in specific hippocampal subfields. Adult animals with a history of postnatal fluoxetine exposure exhibit altered numbers of somatostatin interneurons, GABA receptor subunit expression and neuronal activation within the hippocampus. This indicates that postnatal fluoxetine disrupts the normal developmental trajectory of the hippocampus.

## Introduction

Selective serotonin reuptake inhibitors (SSRIs) are conventionally used as the first line of treatment for women with gestational and postpartum depression (Kodish et al., 2011; Nonacs & Cohen, 2003). Perinatal exposure to SSRIs has been linked to altered neurobehavioral development (Glover & Clinton, 2016; Mulder et al., 2011) and increased risk for suicidal ideation in children and adolescents (Cipriani et al., 2005). Preclinical studies investigating perinatal exposure to SSRIs report increased anxiety- and despair-like behaviors in rodent models that persist across the life-span (Ansorge et al., 2004, 2008; Sarkar et al., 2014). In addition to altering the development of emotionality, perinatal SSRI administration also evokes dysregulation of circadian rhythms and sleep (Popa et al., 2008), alters cortical network function (Simpson et al., 2011), perturbs hormonal stress responses (Norcross et al., 2008), disrupts social play (Bond et al., 2020), reproductive and maternal care behavior (Svirsky et al., 2016), and alters the development of serotonergic circuitry (Weaver et al., 2010). Perinatal SSRI exposure alters hippocampal plasticity (Karpova et al., 2011), neurotrophic factor signaling (Rayen et al., 2015), and disrupts hippocampal neurogenesis in a sexually dimorphic manner (Gemmel et al., 2016). Given the extended developmental trajectory of the hippocampus and the key role it plays in emotional behavior (Rice & Barone, 2000), this limbic brain region that receives dense serotonergic innervation (Oleskevich & Descarries, 1990) is likely to be impacted by perinatal SSRI administration.

The hippocampus exhibits protracted development with neurogenesis, gliogenesis, cell type specification and synaptogenesis extending well into the juvenile window for rodents (Bayer, 1980). Preclinical models of early life stress are reported to evoke a decline in hippocampal neurogenesis (Naninck et al., 2015), dendritic atrophy (Champagne et al., 2008), enhanced cell death (Bath et al., 2016) and altered neurotrophic signaling in the hippocampus (Suri et al., 2014). These structural and functional changes in the hippocampus are implicated in mediating the sustained changes in mood-related behavior that arise as a consequence of early adversity (Chen & Baram, 2016; Fenoglio et al., 2006). The postnatal window marks a critical period in hippocampal development and closure of this critical period is associated with the deposition of extracellular matrix (ECM) moieties, called perineuronal nets (PNNs), preferentially deposited around interneurons (Hensch, 2005). Maternal separation-based models of early stress are reported to cause a decline of PNNs in the prelimbic prefrontal cortex in juvenile animals (Gildawie et al., 2020), and an increase in PNN intensity around Parvalbumin (PV) interneurons in the hippocampus in adults (Murthy et al., 2019). Chronic administration of SSRIs can also impact PNN deposition around PV cells, and is implicated in fear erasure in the basolateral amygdala (Karpova et al., 2011) and in the reopening of critical period plasticity in the visual cortex, likely through the dissolution of PNNs in adulthood (Vetencourt et al., 2008). We hypothesized that SSRI administration during postnatal life could impact the formation and maintenance of PNNs in the hippocampus, a brain region reported to be highly sensitive to perturbations of serotonergic signaling.

In this study, we investigated the influence of postnatal fluoxetine (PNFlx) treatment on the formation of PNNs during postnatal life and their maintenance in adulthood. We addressed whether the number of PNNs that encapsulate PV-positive interneurons in distinct hippocampal subfields are altered as a consequence of PNFlx treatment, both at an early time-point in postnatal life soon after cessation of the SSRI treatment and in adulthood. PNNs are known for the role they play in regulating synaptic physiology and receptor composition, and can thus impinge on neuronal excitation-inhibition balance (Fawcett et al., 2019; Sorg et al., 2016). We also examined the influence of PNFlx treatment on interneuron numbers, NMDA and GABA receptor subunit composition, neuronal activity in the hippocampus and the cytoarchitecture of hippocampal pyramidal neurons. We find that PNFlx treatment results in reduced numbers of PNNs in the hippocampus, and that this decline is maintained into adulthood in a hippocampal subfield-specific manner. This reduction in PNNs results in a significant decline in the number of PV cells ensheathed by PNNs. Additionally, we also observed an increase in neuronal activity within the hippocampus in animals with a history of PNFlx, associated with an altered subunit composition in GABA-A receptors, and a small but significant increase in the dendritic complexity noted in the distal regions of CA1 pyramidal neuron dendritic arbors. Taken together, our results indicate that chronic exposure to SSRIs in the postnatal window alters the trajectory of PNN development, and has implications for the function of local inhibitory circuits in the hippocampus.

## Materials and Methods

### Animals

Sprague–Dawley rats bred at the Tata Institute of Fundamental Research (TIFR) animal facility were used for all experiments. Animals were maintained on a 12-hour light-dark cycle (7 am to 7 pm) with ad libitum access to food and water. Experimental procedures were carried out as per the guidelines of the Committee for the Purpose of Control and Supervision of Experiments on Animals (CPCSEA), Government of India and were approved by the TIFR animal ethics committee (TIFR/IAEC/2017-2).

### Drug Treatments

Litters derived from primiparous Sprague-Dawley dams were assigned at random to either the vehicle or postnatal fluoxetine (PNFlx) administered treatment groups, with each treatment group containing pups obtained from four or more litters, to avoid any litter-specific effects. Rat pups received oral administration of fluoxetine (10 mg/kg; kind gift from IPCA Laboratories) or vehicle (5% sucrose, Sigma-Aldrich, Cat no.) once daily through a micropipette (0.5-10 μL, Eppendorf, 3123000020) from postnatal day 2 (P2) to postnatal day 21 (P21), commencing within 3 minutes of separation from the dam to minimize any handling related stress. Male rat pups were weaned between postnatal days 25-30, following which they were housed in identical group-housing conditions (3 - 4 animals per cage).

### Immunohistochemistry

Sprague-Dawley male rats were sacrificed on postnatal day 21 and postnatal day 80 (adult) via transcardial perfusion with 4% Paraformaldehyde. Harvested brains were sectioned on a VT1000S vibratome (Leica, Germany) to generate serial coronal sections (40 μm thickness). Six free floating hippocampal sections (240 μm apart) per animal were processed for each immunohistochemical staining. Sections were blocked in 10% horse serum made in 0.1M Phosphate Buffer with 0.3% Triton-x (PBTx) for two hours followed by overnight incubation with the following primary antibodies: mouse anti-Parvalbumin (1:1000, Sigma Aldrich, P3088), goat anti-Somatostatin (1:350, Santa Cruz, SC-7819), goat anti-Calretinin (1:1000, Santa Cruz, SC-26512), mouse anti-Reelin (1:1000, Sigma Aldrich, MAB5364) and rabbit anti-c-Fos (1:1000, Cell Signalling Technology, #2250) in 0.3% PBTx, at room temperature. Post washes, the sections were incubated with secondary antibody solutions of biotinylated horse anti-mouse (1:500, Vector Laboratories, BA2000), biotinylated donkey anti-rabbit (1:500, Millipore, AP182B) and biotinylated rabbit anti-goat (1:500, Millipore, AP106.) in 0.3% PBTx for three hours. Subsequently, sections were incubated with Avidin-Biotin complex (Vector Labs, PK-6100) in 0.1M Phosphate Buffer (PB) for one and half hours and visualized with diaminobenzidine staining (Sigma Aldrich, D4293). To detect PNNs. sections were incubated with the plant lectin, biotinylated Wisteria Floribunda (WFA) (1:250, Vector Laboratories, B1355) in 0.3% PBTx overnight followed by incubation with secondary antibody solution of 488 Alexa-Fluor conjugated donkey anti-streptavidin (1:500, Invitrogen, S11223) for 3 hours. For double immunostainings of PV and PNN, sections were incubated with primary antibody solution of rabbit anti-Parvalbumin (1:1000 Abcam UK. ab 11427) and biotinylated Wisteria Floribunda (1:250 Vector Laboratories, B1355) in 0.3% PBTx overnight followed by incubation with secondary antibody solutions of 488 Alexa-Fluor conjugated donkey anti-streptavidin (1:500, Invitrogen USA, S11223) and 555 Alexa-Fluor conjugated donkey anti-rabbit (1:500, Invitrogen, A31572) for 3 hours.

### Cell visualization and counting

Immunostained sections were mounted on glass slides with DPX mountant medium (Merck, 100579). Slides were coded and quantification was carried out by the experimenter blind to the treatment groups. Cells were visualised at 20X magnification using a bright field microscope (Zeiss Axioscope 2, Germany). Total number of stained cells were counted per animal per hippocampal subfield, namely the CA1, CA3 and DG, and divided by the number of sections to obtain an average number of cells per section within the respective hippocampal subfield. Results for immunostaining were expressed as mean ± SEM. For PV and PNN double immunostainings, sections from each animal were mounted on glass slides with Vectashield Hard-set Antifade mounting media with DAPI (Vector Labs, H-1500). Slides were coded and quantification was carried out by the experimenter blind to the treatment groups. Cells were visualised at 20X magnification and images were acquired using the Zeiss Axio Imager M2 (Zeiss, Germany). The percent colocalization of PV-positive cells with the PNN marker WFA was determined in the DG, CA1, CA3 hippocampal subfield in six sections (250 μm apart) per animal. A minimum of 50 PV-positive cells were analyzed per animal using Z-plane sectioning (0.5 μm) to confirm the percent colocalization of PV with the PNN marker WFA.

### Quantitative polymerase chain reaction

Quantitative PCR (qPCR) was performed on Vehicle and PNFlx treatment groups at two time points, namely P21, 2 hours after cessation of the final fluoxetine/vehicle treatment and in adulthood (postnatal day 100). The hippocampus was micro-dissected, snap-frozen using liquid nitrogen, and kept at −80°C for long-term storage. RNA was extracted using TRI reagent (Sigma-Aldrich Chemicals ltd.) and reverse transcribed using complementary DNA (cDNA) synthesis kit (PrimeScript 1st strand cDNA Synthesis Kit, Takara Bio). The synthesized cDNA was subjected to qPCR using KAPA SYBR (KAPABiosystems) and a Bio-Rad CFX96 real-time PCR machine. Primers were designed using the NCBI primer BLAST program. The complete list of primers used is provided in Table 1. Hypoxanthine Phosphoribosyltransferase 1 (*Hprt-1*) was used as the housekeeping gene for normalization.

**Table 1.**
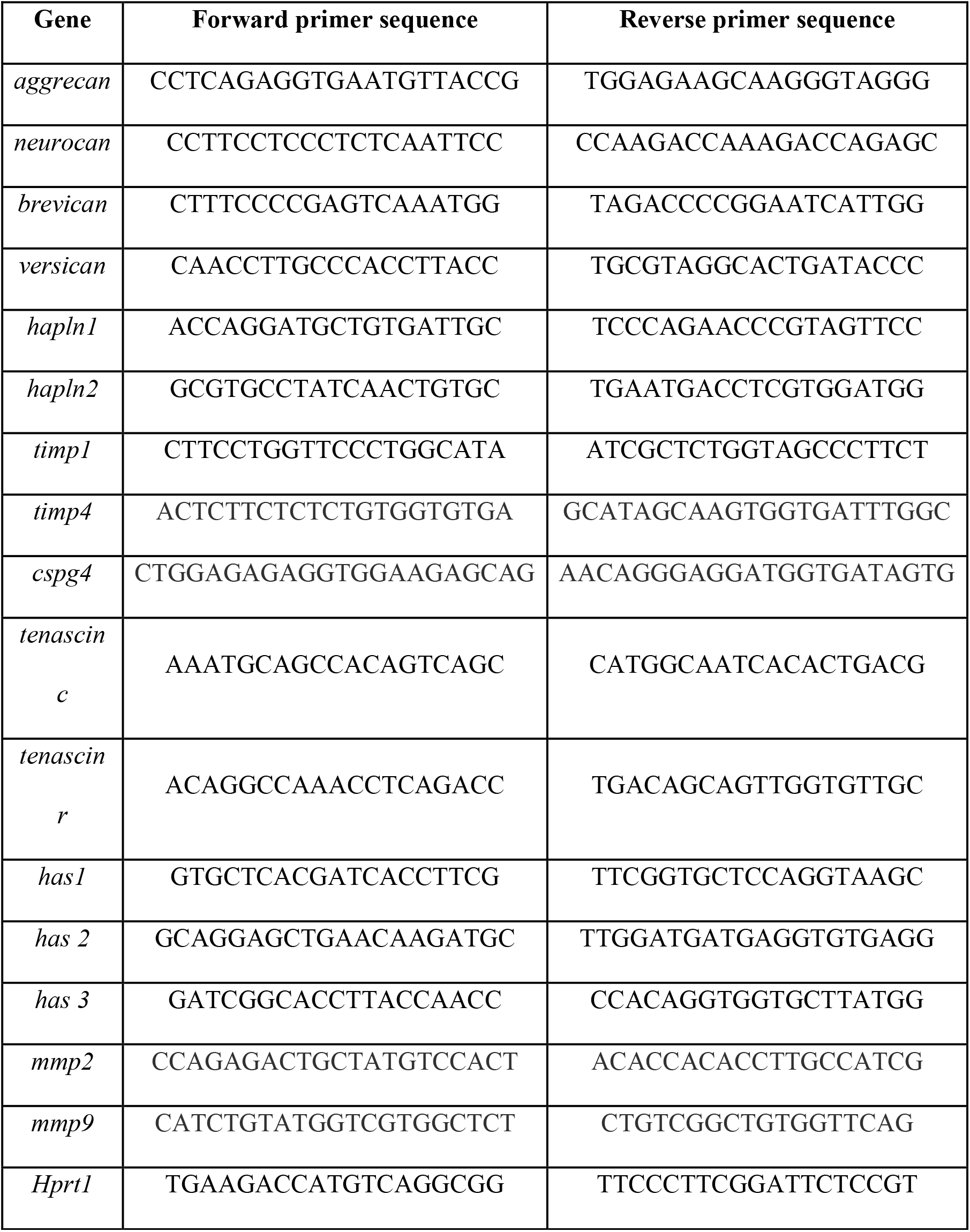
Primer sequences (5’-3’)

### Western Blotting

Male Sprague-Dawley rats were sacrificed on postnatal day 100 and their hippocampi were dissected, snap-frozen in liquid nitrogen and stored at −80°C. Tissue homogenization and protein extraction was performed in radioimmunoprecipitation assay (RIPA) buffer (10 mM Tris-Cl (pH 8.0), 1 mM EDTA, 0.5 mM EGTA, 1% Triton X-100, 0.1% sodium deoxycholate, 0.1% SDS, 140 mM NaCl). Protease inhibitor cocktail (Sigma Aldrich, P8340), phosphatase inhibitor cocktail 2 (Sigma-Aldrich, P5276), and phosphatase inhibitor cocktail 3 (Sigma Aldrich, P0044) were added to the extraction buffer prior to protein extraction. Protein concentrations were estimated using a Quantipro BCA assay kit (Sigma Alrich). Protein lysates (50 μg) were resolved on a sodium dodecyl sulphate polyacrylamide gel electrophoresis (SDS-PAGE) system and transferred onto polyvinylidene fluoride membranes. The membranes were blocked using 5% bovine serum albumin (Sigma-Aldrich, A9418) in Tris-buffered saline with 0.1% of Tween-20 (TBST) followed by incubation with the primary antibodies, namely rabbit anti-NR2A (1:1000, Millipore, 07-632), rabbit anti-NR2B (1:1000, Millipore, 06-600), rabbit anti-GABA-Aa1 (1:1000, Abcam, ab33299), mouse anti-GABA-Aa2 (1:1000, Abcam, ab193311), and rabbit anti-ß-Actin (1:10000, Abclonal, AC026) in TBST with 5% Bovine Serum Albumin (BSA) overnight at 4°C. Following washes, the blots were incubated with horseradish peroxidase conjugated goat anti-rabbit (1:5000, Abclonal, AS014) or horseradish peroxidase conjugated goat anti-mouse (1:5000, Abclonal, AS003) for two hours. Visualization of the signal was carried out using a western blotting detection kit (SuperSignal West Pico Plus, Thermofisher, 34579) with the GE Amersham Imager 600 (GE life sciences). Densitometric analysis of the blots were performed using ImageJ (National Institute of Health, USA).

### Golgi staining and arborization analysis

Male Sprague-Dawley rats were sacrificed at postnatal day 120 and their brains were harvested and Golgi staining was carried out using the FD Rapid GolgiStain™ Kit (PK401). Each brain was cut coronally into a smaller chunk containing the entire hippocampus and incubated with the impregnation solution (solution A and B) in the dark for 21 days. After impregnation, the brains are placed in solution C for 3 days. Sections were taken on a vibratome (Leica, Wetzlar, Germany) at a thickness of 150 μm, and incubated with freshly prepared staining solutions as per the instructions in the FDRapid GolgiStain™ Kit for 10 mins. Post washes, the sections on slides were dehydrated using xylene, and mounted with DPX mountant medium (Merck, 100579). For tracing of neurons, slides were coded and quantification was carried out by the experimenter blind to the treatment groups. Tracing of CA1 pyramidal neurons was carried out at 20X magnification on the BX53 light microscope (Olympus, Japan) using the Neurolucida 10 (MBF Biosciences, USA). For analysis of traces, the Neurolucida 10 Explorer (MBF Biosciences, USA) was used to perform Sholl analysis.

### Statistical Analysis

All experiments had two treatment groups and were subjected to the Shapiro-Wilk test to determine normality of the data. Data that followed a normal distribution were subjected to a two-tailed, unpaired Student’s t-test using GraphPad Prism 8 (Graphpad Software Inc., USA). Data that did not exhibit a normal distribution pattern were subjected to the Mann-Whitney U test. Graphs were plotted using GraphPad Prism (Graphpad Software Inc., USA). Data are expressed as mean ± standard error of the mean (S.E.M) and statistical significance was set at *p* < 0.05. Welch’s correction was applied when variances varied significantly between the treatment groups. For the arborization analysis, Friedman's test of repeated measures for non-parametric distributions was performed for Sholl analysis and results were considered significant when *p* < 0.05. A detailed statistical summary for all figures is provided in Table 2 and Table 3.

**Table 2.** Statistical table for figures.

| Figure number | Figure detail | Data Structure | Type of test | Confidence interval | *p* value |
| --- | --- | --- | --- | --- | --- |
| 1 | B | Normal | Student's *t*-test | 21.30 to 53.30 | ***0.0003*** |
|  | C | Normal | Student's *t*-test | 4.40 to 15.40 | ***0.0023*** |
|  | D | Normal | Student's *t*-test | -4.73 to 1.23 | ***0.2002*** |
|  | F; PV<sup>+</sup> WFA<sup>-</sup> | Non-normal | Mann-Whitney U Test | -0.46 to -0.05 | ***0.0286*** |
|  | F; PV<sup>+</sup> WFA<sup>+</sup> | Non-normal | Mann-Whitney U Test | 0.05 to 0.46 | ***0.0286*** |
|  | G; PV<sup>+</sup> WFA<sup>-</sup> | Non-normal | Mann-Whitney U Test | -0.43 to -0.09 | ***0.0286*** |
|  | G; PV<sup>+</sup> WFA<sup>+</sup> | Non-normal | Mann-Whitney U Test | 0.09 to 0.43 | ***0.0286*** |
|  | H; PV<sup>+</sup> WFA<sup>-</sup> | Normal | Student's *t*-test | -0.13 to 0.03 | *0.1990* |
|  | H; PV<sup>+</sup> WFA<sup>+</sup> | Normal | Student's *t*-test | -0.03 to 0.13 | *0.1990* |
| 2 | B | Normal | Student's *t*-test | -23.71 to -0.78 | ***0.0379*** |
|  | C | Normal | Student's *t*-test | -9.31 to 0.79 | *0.0922* |
|  | D | Normal | Student's *t*-test | -5.38 to 4.70 | *0.8863* |
|  | F; PV<sup>+</sup> WFA<sup>-</sup> | Normal | Student's *t*-test | -0.08 to 0.05 | *0.6638* |
|  | F; PV<sup>+</sup> WFA<sup>+</sup> | Normal | Student's *t*-test | -0.05 to 0.08 | *0.6638* |
|  | G; PV<sup>+</sup> WFA<sup>-</sup> | Normal | Student's *t*-test | -0.14 to 0.10 | *0.5950* |
|  | G; PV<sup>+</sup> WFA<sup>+</sup> | Normal | Student's *t*-test | -0.10 to 0.14 | *0.5950* |
|  | H; PV<sup>+</sup> WFA<sup>-</sup> | Non-normal | Mann-Whitney U Test | -0.10 to 0.16 | *0.6870* |
|  | H; PV<sup>+</sup> WFA<sup>+</sup> | Non-normal | Mann-Whitney U Test | -0.16 to 0.10 | *0.6870* |
| 4 | C; CA1 | Normal | Student's *t*-test | -85.40 to 24.43 | *0.2306* |
|  | C; CA3 | Normal | Student's *t*-test | -63.14 to 12.67 | *0.1594* |
|  | C; DG | Normal | Student's *t*-test | -16.70 to 12.98 | *0.7698* |
|  | D; CA1 | Normal | Student's *t*-test | -40.69 to 24.94 | *0.5785* |
|  | D; CA3 | Normal | Student's *t*-test | -10.45 to 23.48 | *0.3836* |
|  | D; DG | Normal | Student's *t*-test | -7.07 to -3.43 | ***0.0007*** |
|  | F; CA1 | Normal | Student's *t*-test | -21.39 to 23.18 | *0.9284* |
|  | F; CA3 | Normal | Student's *t*-test | -12.14 to 36.55 | *0.2811* |
|  | F; DG | Non-normal | Mann-Whitney U Test | -15.03 to 20.72 | *0.4206* |
|  | G; CA1 | Non-normal | Mann-Whitney U Test | -32.39 to 98.72 | *0.6623* |
|  | G; CA3 | Normal | Student's *t*-test | -67.94 to 42.92 | *0.6220* |
|  | G; DG | Non-normal | Mann-Whitney U Test | -31.82 to 7.71 | *0.2468* |
|  | I; CA1 | Normal | Student's *t*-test | -78.79 to 33.99 | *0.3321* |
|  | I; CA3 | Normal | Student's *t*-test | -26.61 to 3.01 | *0.0914* |
|  | I; DG | Normal | Student's *t*-test | -10.22 to 20.88 | *0.3950* |
|  | J; CA1 | Normal | Student's *t*-test | -34.27 to 6.37 | *0.1549* |
|  | J; CA3 | Normal | Student's *t*-test | -10.45 to 23.48 | *0.6372* |
|  | J; DG | Normal | Student's *t*-test | -17.16 to 14.47 | *0.8533* |
|  | L; CA1 | Normal | Student's *t*-test | -79.33 to 41.76 | *0.5022* |
|  | L; CA3 | Normal | Student's *t*-test | -14.10 to 19.65 | *0.7234* |
|  | L; DG | Normal | Student's *t*-test | -31.34 to 24.28 | *0.7818* |
|  | M; CA1 | Normal | Student's *t*-test | -54.22 to 10.70 | *0.1725* |
|  | M; CA3 | Normal | Student's *t*-test | -14.59 to 3.37 | *0.2017* |
|  | M; DG | Normal | Student's *t*-test | -10.77 to 8.72 | *0.8242* |
| 5 | C; NR2A | Normal | Student's *t*-test | -0.83 to 1.50 | *0.4880* |
|  | C; NR2B | Normal | Student's *t*-test | -1.14 to 1.03 | *0.9083* |
| | E; GABA-Aα1 | Normal | Student's *t*-test | -0.06 to 0.94 | *0.0723* |
| | E; GABA-Aα2 | Normal | Student's *t*-test | -0.83 to -0.05 | ***0.0330*** |
|  | G; CA1 | Non-normal | Mann-Whitney U Test | 2.095 to 27.55 | ***0.0175*** |
|  | G; CA3 | Non-normal | Mann-Whitney U Test | 1.60 to 8.69 | ***0.0105*** |
|  | G; DG | Normal | Student's *t*-test | -46.91 to -5.60 | ***0.0170*** |
|  | I | Non-normal | Friedman's test |  | ***<0.001*** |
|  | J | Non-normal | Friedman's test |  | ***0.0170*** |

**Table 3:** Statistical table for Figure 3.

| Gene | P21 |  |  |  | Adult |  |  |  |
| --- | --- | --- | --- | --- | --- | --- | --- | --- |
|  | Data Structure | Type of test | Confidence Interval | *p* value | Data Structure | Type of test | Confidence Interval | *p* value |
| *aggrecan* | Normal | Student's t-test | -0.45 to 0.36 | *0.8008* | Normal | Student's t-test | -0.43 to 0.03 | *0.0881* |
| *neurocan* | Normal | Student's t-test | -0.72 to 0.13 | *0.1531* | Non-normal | Mann-Whitney U test | -0.34 to 0.94 | *0.3702* |
| *brevican* | Normal | Student's t-test | -0.73 to 0.29 | *0.3640* | Normal | Student's t-test | -0.17 to 0.65 | *0.2335* |
| *versican* | Normal | Student's t-test | -0.64 to 0.09 | *0.1301* | Normal | Student's t-test | -0.20 to 0.10 | *0.4973* |
| *hapln1* | Non-normal | Mann-Whitney U test | -0.54 to 0.21 | *0.8357* | Normal | Student's t-test | -0.22 to -0.06 | *0.2205* |
| *hapln2* | Normal | Student's t-test | -0.60 to 0.26 | *0.4059* | Normal | Student's t-test | 0.35 to 0.86 | *0.1137* |
| *timp1* | Normal | Student's t-test | -0.37 to 0.19 | *0.5017* | Normal | Student's t-test | -0.04 to 0.26 | *0.6473* |
| *timp4* | Non-normal | Mann-Whitney U test | -0.41 to 0.17 | *0.9999* | Normal | Student's t-test | -0.15 to 0.54 | *0.7113* |
| *cspg4* | Normal | Student's t-test | -0.39 to 0.21 | *0.5284* | Non-normal | Mann-Whitney test | -0.16 to 0.92 | *0.2799* |
| *tenascin c* | Non-normal | Mann-Whitney U test | -0.48 to 0.37 | *0.9452* | Normal | Student's t-test | -0.32 to -0.20 | ***0.0054*** |
| *tenascin r* | Non-normal | Mann-Whitney U test | -1.04 to 0.09 | *0.1014* | Normal | Student's t-test | -0.93 to 0.42 | *0.4397* |
| *has1* | Normal | Student's t-test | -0.35 to 0.40 | *0.8841* | Normal | Student's t-test | -0.30 to -0.10 | *0.1810* |
| *has 2* | Non-normal | Mann-Whitney U test | -0.30 to 0.20 | *0.6282* | Normal | Student's t-test | -0.30 to -0.09 | *0.2027* |
| *has 3* | Normal | Student's t-test | -0.40 to 0.19 | *0.4629* | Normal | Student's t-test | -0.40 to -0.24 | ***0.0129*** |
| *mmp2* | Normal | Student's t-test | -0.60 to -0.004 | ***0.0474*** | Non-normal | Mann-Whitney U test | -0.69 to 1.04 | *0.1564* |
| *mmp9* | Normal | Student's t-test | -0.38 to 0.03 | *0.0881* | Non-normal | Mann-Whitney U test | -0.43 to 0.50 | *0.9742* |

## Results

### 1.3.1 Postnatal fluoxetine treatment alters perineuronal net numbers in the hippocampus

PNNs are extracellular matrix (ECM) based structures that play a key role in the maturation of neural circuits across development, and are known to exhibit substantial alterations in response to environmental perturbations during critical period windows (Sorg et al., 2016). To study the effects of postnatal fluoxetine (PNFlx) exposure on PNNs in the hippocampal subfields, we orally administered fluoxetine to neonatal rats from postnatal day 2 to 21 (P2-P21) (Fig 1A, Fig 2A). At P21 and adulthood, we examined the expression of Wisteria Floribunda (WFA), a plant lectin that binds to the sugar moieties present on the PNNs (Brückner et al., 1996). This allowed us to address the short- and long-term influence of elevation of serotonin levels during the postnatal temporal window via PNFlx treatment on both the formation and maintenance of PNNs in the hippocampus.

**Figure 1.**
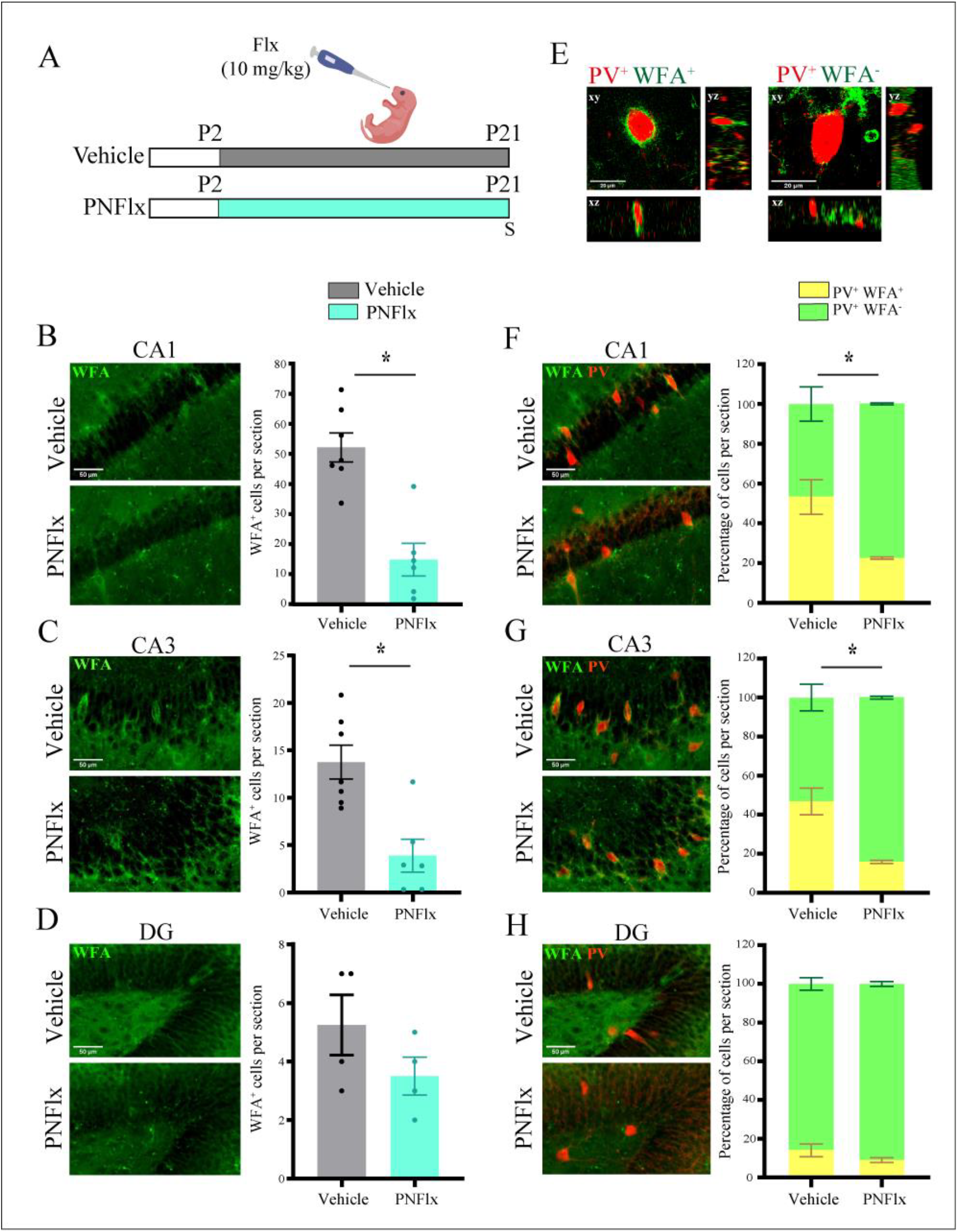
Influence of postnatal fluoxetine treatment on perineuronal nets in the hippocampus at P21. (A) Shown is a schematic depicting the experimental paradigm used to test the effects of postnatal fluoxetine (PNFlx) administration from P2 to P21 on hippocampal PNNs. WFA staining was performed to detect PNNs and to assess the number of PV-positive neurons that were surrounded by PNNs. S denotes the age (P21) at which animals were sacrificed. (B) Shown are representative images and quantification of WFA-positive PNNs in the CA1 hippocampal subfields from vehicle and PNFlx-treated animals at P21. PNNs in the CA1 hippocampal subfields of PNFlx animals showed a significant decline compared to vehicle treated animals. (C) Shown are representative images and quantification of WFA-positive PNNs in the CA3 hippocampal subfields from vehicle and PNFlx treated animals at P21. PNNs in the CA3 hippocampal subfields of PNFlx animals showeda significant decline compared to vehicle treated animals. (D) Shown are representative images and quantification of WFA-positive PNNs in the DG hippocampal subfields from vehicle and PNFlx-treated animals at P21. No significant difference was seen in the number of PNNs in the DG hippocampal subfields of vehicle and PNFlx-treated animals at P21. (E) Shown are representative confocal z-stacks of PV-positive neurons which exhibit the presence (left) or absence (right) of co-localization with a WFA-stained PNN. (F) Shown are representative double immunofluorescence images of WFA-positive PNNs (green) and PV-positive (red) neurons in the CA1 hippocampal subfields from vehicle and PNFlx-treated animals at P21. Significantly lesser percent of PV-positive cells are surrounded by PNNs in the CA1 hippocampal subfields of PNFlx animals compared to vehicle treated animals. (G) Shown are representative double immunofluorescence images of WFA-positive PNNs (green) and PV-positive (red) neurons in the CA3 hippocampal subfields from vehicle and PNFlx-treated animals at P21. Significantly lesser percent of PV-positive cells are surrounded by PNNs in the CA3 hippocampal subfields of PNFlx animals compared to vehicle treated animals. (H) Shown are representative double immunofluorescence images and quantification of WFA-positive PNNs (green) and PV-positive (red) neurons in the DG hippocampal subfields from vehicle and PNFlx-treated animals at P21. Quantification revealed no significant difference between the DG hippocampal subfields of vehicle and PNFlx-treated animals at P21. Results are expressed as the mean ± SEM, \**p*<0.05 as compared to vehicle treated control animals, two-tailed unpaired Student’s t-test or Mann-Whitney U test.

**Figure 2.**
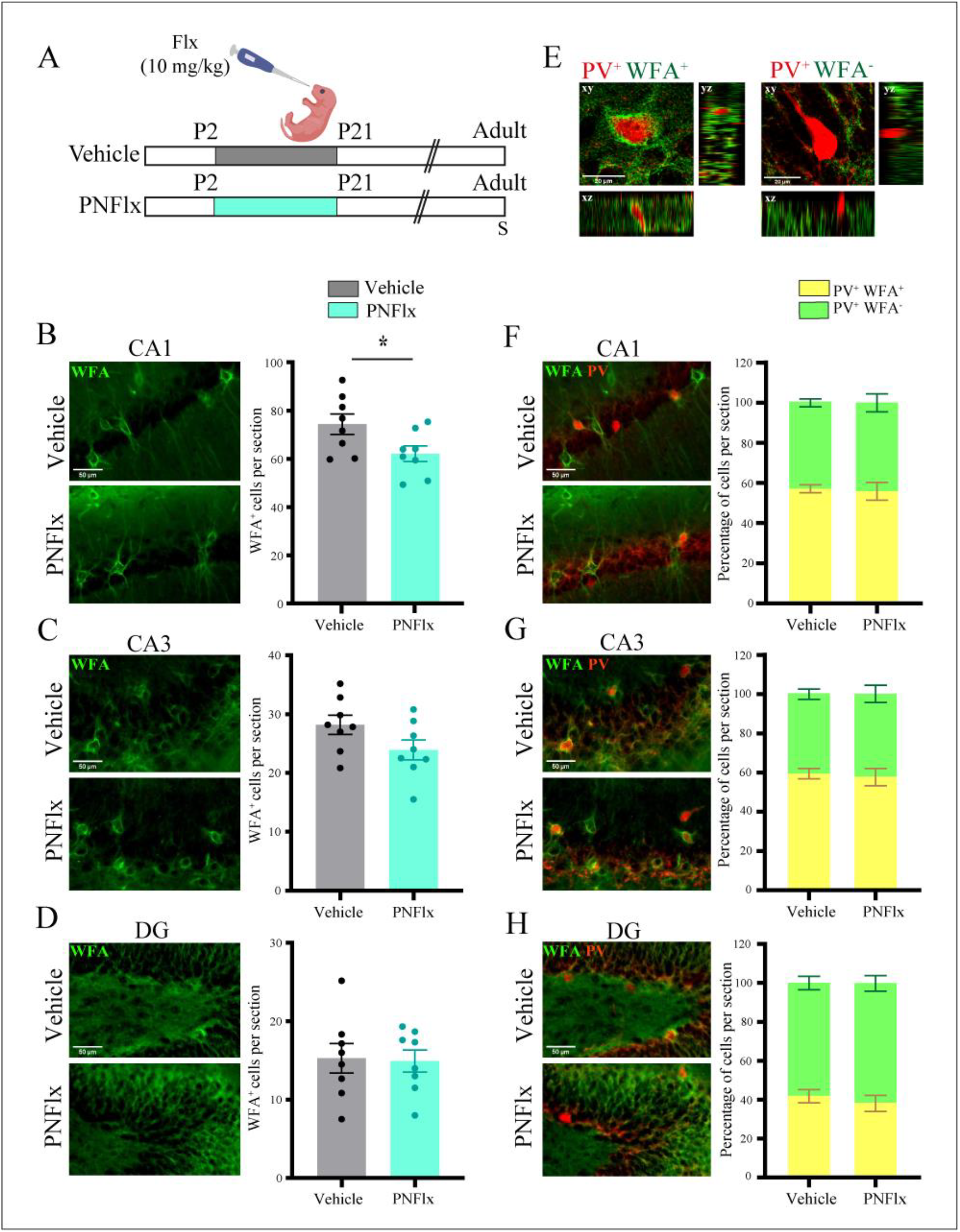
Influence of postnatal fluoxetine treatment on perineuronal nets in the hippocampus in adulthood. (A) Shown is a schematic depicting the experimental paradigm used to test the effects of PNFlx administration from P2 to P21 on hippocampal PNNs. WFA staining was performed to detect PNNs and to assess the number of PV-positive neurons that were surrounded by PNNs. S denotes the age (P80) at which animals were sacrificed. (B) Shown are representative images and quantification of WFA-positive PNNs in the CA1 hippocampal subfields from vehicle and PNFlx-treated adult animals. PNNs in the CA1 hippocampal subfields of PNFlx animals showed a significant decline compared to vehicle treated animals. (C) Shown are representative images and quantification of WFA-positive PNNs in the CA3 hippocampal subfields from vehicle and PNFlx-treated adult animals. No significant difference was seen in the number of PNNs in the CA3 hippocampal subfields of vehicle and PNFlx-treated animals in adulthood. (D) Shown are representative images and quantification of WFA-positive PNNs in the DG hippocampal subfields from PNFlx and vehicle treated adult animals. No significant difference was seen in the number of PNNs in the DG hippocampal subfields of vehicle and PNFlx-treated animals in adulthood. (E) Shown are representative confocal z-stacks of PV-positive neurons which exhibit the presence (left) or absence (right) of co-localization with a WFA-stained PNN. (F) Shown are representative double immunofluorescence images and quantification of WFA-positive PNNs (green) and PV-positive (red) neurons in the CA1 hippocampal subfields from vehicle and PNFlx-treated adult animals. (G) Shown are representative double immunofluorescence images and quantification of WFA-positive PNNs (green) and PV-positive (red) neurons in the CA3 hippocampal subfields from vehicle and PNFlx-treated adult animals. (H) Shown are representative double immunofluorescence images and quantification of WFA-positive PNNs (green) and PV-positive (red) neurons in the DG hippocampal subfields from vehicle and PNFlx-treated adult animals. Quantification revealed no significant difference between theCA1, CA3 and DG hippocampal subfields of vehicle and PNFlx-treated animals in adulthood. Results are expressed as the mean ± SEM, \**p*<0.05 as compared to vehicle treated control animals, two-tailed unpaired Student’s *t-*test or Mann-Whitney U test.

At P21, we observed a robust decline in the number of WFA-stained PNNs in the CA1 (Fig 1B; *p* value = 0.0003, n = 6 - 7 per group) and CA3 (Fig 1C) (*p* value = 0.002, n = 6 - 7 per group) subfields of the hippocampus in PNFlx animals compared to the vehicle-treated control group. In contrast, we noted no difference in the number of PNNs detected within the DG subfield (Fig 1D). As temporal windows of critical period plasticity close, PNNs condense primarily around local GABAergic PV-positive interneurons across numerous cortical and subcortical regions, restricting extensive experience-dependent remodeling within these circuits (Hensch, 2004). We next analyzed whether PNFlx treatment influenced the fraction of hippocampal PV-positive interneurons that were surrounded by PNNs (Fig 1E) at the early time point of P21. We find a significantly smaller fraction of PV-positive neurons surrounded by PNNs in the CA1 (Fig 1F; PNN/PV_Vehicle/CA1_ = 53.25%, PNN/PV_PNFlx/CA1_ = 23.25%; *p* value = 0.029, n = 4 per group) and CA3 (Fig 1G; PNN/PV_Vehicle/CA3_ = 46.75%, PNN/PV_PNFlx/CA3_ = 15.75%; *p* value = 0.029, n = 4 per group) subfields of PNFlx-treated animals than in the vehicle-treated controls. Within the DG subfield, we noted no change in the colocalization of PV-positive neurons with WFA-positive PNNs as a consequence of PNFlx treatment (Fig 1H). Our results indicate that there are marked changes in the total numbers of PNNs, as well as in the colocalization of PV-positive neurons with PNNs, within specific hippocampal subfields immediately after the cessation of PNFlx treatment.

We next sought to address whether the decline in PNN numbers noted in specific hippocampal subfields with PNFlx treatment at P21 persists into adulthood. We find that the CA1 hippocampal subfield of PNFlx animals continued to exhibit a deficit in the total number of PNNs (Fig 2B; *p* value = 0.038, n = 8 per group). In contrast, the number of PNNs in the CA3 hippocampal subfield in the PNFlx animals did not differ from vehicle-treated controls, indicating that the decline noted in PNN numbers in this subfield at P21 had shown substantial recovery by adulthood (Fig 2C). There was no difference in the number of PNNs in the DG hippocampal subfield in adulthood in the PNFlx cohort (Fig 2D). We did not observe a difference in the proportion of PV-positive neurons that were surrounded by PNNs in either the CA1, CA3 or DG subfield (Fig 2F-H). We note that the WFA-stained PNNs in adulthood (Fig 2E) were better formed and showed a more prominent structure as compared to those at P21 (Fig 1E). This is consistent with age-dependent maturation of the PNN structure that has been reported in many areas of the brain previously (Lipachev et al., 2019; Pizzorusso et al., 2002). Our results suggest that a transient elevation of serotonin in the postnatal temporal window alters the time-course of the appearance and maturation of PNNs around PV-positive neurons in a subfield-specific manner in the hippocampus.

### 1.3.2 Postnatal fluoxetine treatment alters perineuronal net associated gene expression in the hippocampus

We next addressed whether the transcription of PNN components or molecular machinery that is known to enzymatically modulate the ECM were specifically altered as a consequence of PNFlx treatment. We performed qPCR analysis on hippocampal lysates derived from PNFlx and vehicle-treated animals at P21 and in adulthood to examine potential short- and long-term changes in the expression of different genes associated with the extracellular matrix composition, synthesis, proteolytic breakdown and PNN-linked plasticity (Fig 3A-B). We did not observe any dysregulation in the constituent proteoglycans of PNNs such as *aggrecan*, *brevican*, *neurocan*, *versican* and *cspg4* in the hippocampus at either age following PNFlx treatment. However, at P21, we observe a downregulation of *mmp2* mRNA (*p* value = 0.047, n = 6 - 7 per group), a matrix metalloprotease whose role in dendritic remodelling, synaptic plasticity, and axonal sprouting in the hippocampus has been extensively studied (Fujioka et al., 2012). In adulthood, we observed significant downregulation in the gene expression of *tenascin c*, a glycoprotein, implicated in hippocampal plasticity (Evers et al., 2002) (*p* value = 0.005, n = 8 - 10 per group). *Hyaluronan synthase 3* (Has3), a membrane enzyme involved in the synthesis of a critical PNN ECM component, hyaluronan (Fawcett et al., 2019), was also downregulated in the hippocampi of PNFlx animals in adulthood (*p* value = 0.012, n = 8 - 10 per group). Taken together, our data suggest that while PNFlx treatment adversely affects hippocampal PNNs in both a short- and long-term manner, there is a limited regulation noted of PNN-associated components at the transcriptional level at these time-points.

**Figure 3.**
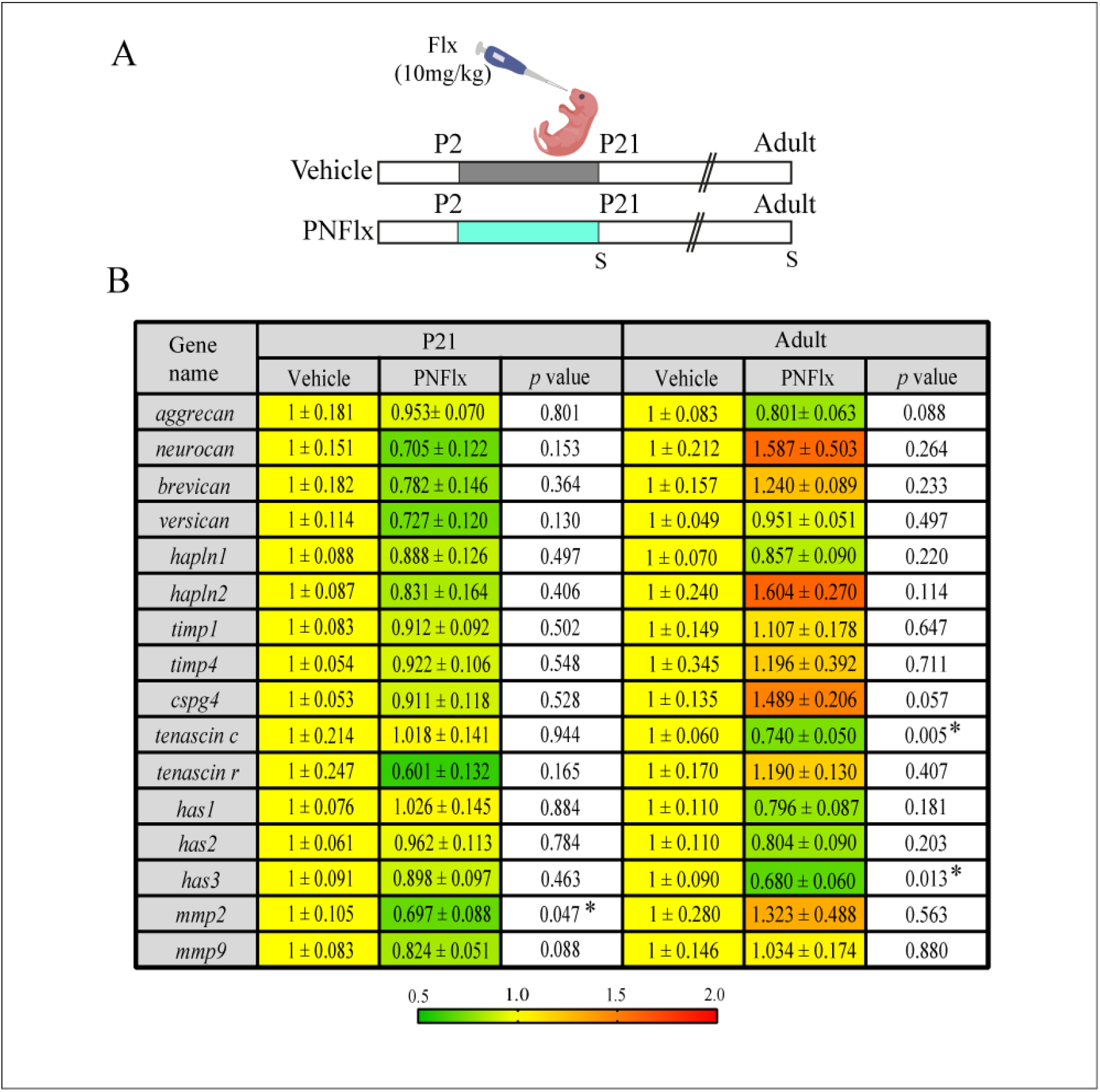
Transcriptional changes of extracellular matrix components in adult animals with a history of postnatal administration of fluoxetine. (A) Shown is a schematic depicting the experimental paradigm used to test the effects of PNFlx administration from P2 to P21 on expression of PNN associated components in the hippocampus of animals at P21 and adulthood. S denotes the ages (P21 and P100) at which animals were sacrificed. (B) Shown are the normalized gene expression levels expressed as fold change relative to their age-matched vehicle treated animals, as well as p-values calculated using two-tailed, unpaired Student’s *t-*test or Mann-Whitney U test (n = 6-11 per group). Heat map indicates the extent of gene regulation. Results are expressed as mean ± SEM. (*hapln1* - hyaluronan and proteoglycan link protein 1, *hapln2* - hyaluronan and proteoglycan link protein 2, *timp1* - tissue inhibitor of metalloproteinases 1, *timp4* - tissue inhibitor of metalloproteinases 4, *cspg4* - chondroitin sulphate proteoglycan 4, *has1* - hyaluronan synthase 1, *has2* - hyaluronan synthase 2, *has3* - hyaluronan synthase 3, *mmp2* - matrix metalloproteinase 2, *mmp9* - matrix metalloproteinase 9).

### 1.3.2 Postnatal fluoxetine does not alter interneuron numbers in the hippocampus

GABAergic interneurons continue to migrate, integrate into circuits, and undergo apoptosis during early postnatal life (Wonders & Anderson, 2006). This temporal window overlaps with the period of PNFlx administration. Interneurons are known to express 5HT3A receptors and the *in vitro* migratory patterns of embryonic interneurons have been reported to be affected by serotonergic signaling (Puig & Gulledge, 2011; Riccio et al., 2009). We thus aimed to investigate the effects of PNFlx treatment on hippocampal interneuron numbers at P21 and adulthood. We performed immunostaining for four major, non-overlapping subclasses of interneurons, that are immunoreactive for Parvalbumin (PV), Somatostatin (SST), Calretinin (CR) and Reelin.

At P21, we observed no significant difference in the number of PV-positive, SST-positive, CR-positive, and Reelin-positive interneurons in the hippocampi of PNFlx-treated animals compared to vehicle-treated controls (Fig 4C, F, I, L). Given we noted a significant decline in the number of PV-positive neurons surrounded by PNNs at P21 in the CA1 hippocampal subfield, this indicates that PNFlx impacts the formation of PNNs around PV-positive neurons in the hippocampus of PNFlx animals, but does not influence total PV-positive neuron number. Similarly, at P80, the numbers of PV-positive, SST-positive, CR-positive and Reelin-positive interneurons present in the CA1, CA3 and DG hippocampal subfields (Fig 4D, G, J, M) was not altered by PNFlx treatment. However, we did observe a small but significant increase in the number of SST-positive neurons in the DG hippocampal subfield of adult animals with a history of PNFlx treatment (*p* value = 0.0007, n = 3 - 4 per group).

**Figure 4.**
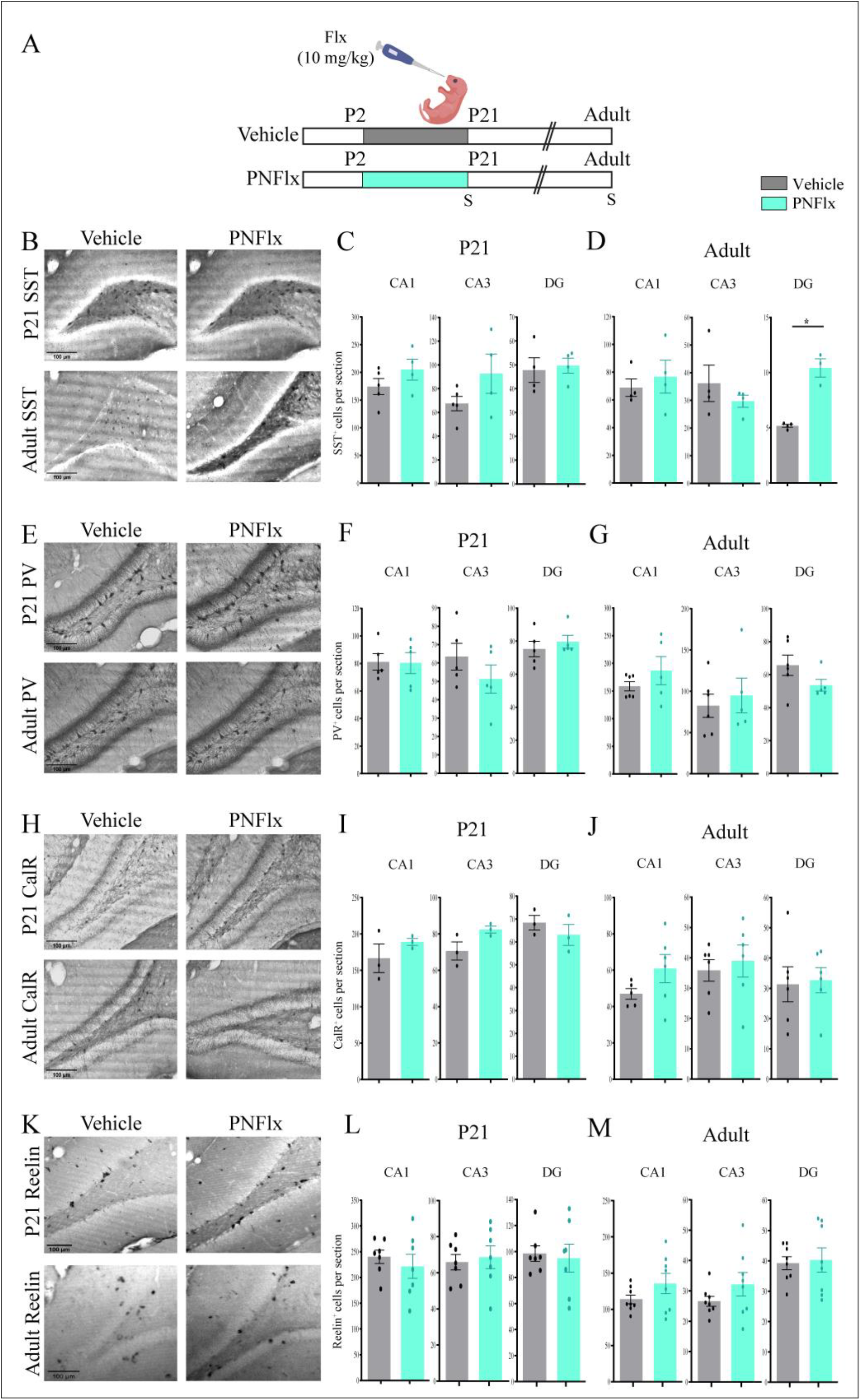
Influence of postnatal fluoxetine administration on interneuron numbers in the postnatal and adult hippocampus. (A) Shown is a schematic depicting the experimental paradigm to test effects of PNFlx administration from P2 to P21 on hippocampal interneurons at P21 and adulthood. S denotes the ages (P21 and P80) at which animals were sacrificed. (B) Shown are representative images of Somatostatin (SST)-positive neurons in the hippocampus of PNFlx- and vehicle-treated animals at P21 and in adulthood. (C) Shown is quantification of SST-positive neurons in theCA1, CA3 and DG hippocampal subfields in PNFlx- and vehicle-treated control animals at P21. Quantification of SST-positive neurons in the CA1, CA3 and DG hippocampal subfields didn’t show a significant change between PNFlx- and vehicle-treated animals at P21. (D) Shown is quantification of SST-positive neurons in the CA1, CA3 and DG hippocampal subfields in PNFlx- and vehicle-treated animals in adulthood. Quantification of SST-positive neurons showed a significant increase in the DG hippocampal subfield of adult PNFlx-treated animals while no change was observed in CA1 and CA3 hippocampal subfields. (E) Shown are representative images of PV-positive neurons in the hippocampus of PNFlx-and vehicle-treated animals at P21 and in adulthood. (F) Shown is quantification of PV-positive neurons in the CA1, CA3 and DG hippocampal subfields in PNFlx- and vehicle-treated animals at P21. Quantification of PV-positive neurons in the CA1, CA3 and DG hippocampal subfields didn’t show a significant change between PNFlx- and vehicle-treated animals at P21. (G) Shown is quantification of PV-positive neurons in the CA1, CA3 and DG hippocampal subfields in PNFlx- and vehicle-treated animals in adulthood. Quantification of PV-positive neurons in the CA1, CA3 and DG hippocampal subfields didn’t show a significant change between PNFlx- and vehicle-treated animals in adulthood. (H) Shown are representative images of Calretinin (CalR)-positive neurons in the hippocampus of PNFlx- and vehicle-treated animals at P21 and in adulthood. (I) Shown is quantification of CalR-positive neurons in the CA1, CA3 and DG hippocampal subfields in PNFlx- and vehicle-treated animals at P21. Quantification of CalR-positive neurons in the CA1, CA3 and DG hippocampal subfields didn’t show a significant change between PNFlx- and vehicle-treated animals at P21. (J) Shown is quantification of CalR-positive neurons in the CA1, CA3 and DG hippocampal subfields in PNFlx- and vehicle-treated animals in adulthood. Quantification of CalR-positive neurons in the CA1, CA3 and DG hippocampal subfields didn’t show a significant change between PNFlx- and vehicle-treated animals in adulthood. (K) Shown are representative images of Reelin-positive neurons in the hippocampus of PNFlx- and vehicle-treated animals at P21 and in adulthood. (L) Shown is quantification of Reelin-positive neurons in the CA1, CA3 and DG hippocampal subfields in PNFlx- and vehicle-treated animals at P21. Quantification of Reelin-positive neurons in the CA1, CA3 and DG hippocampal subfields didn’t show a significant change between PNFlx- and vehicle-treated animals at P21. (M) Shown is quantification of Reelin-positive neurons in the CA1, CA3 and DG hippocampal subfields in PNFlx- and vehicle-treated animals at adulthood. Quantification of Reelin-positive neurons in the CA1, CA3 and DG hippocampal subfields didn’t show a significant change between PNFlx- and vehicle-treated animals in adulthood. All results are expressed as the mean ± SEM, \**p*<0.05 as compared to vehicle treated animals, two-tailed unpaired Student’s t-test or Mann-Whitney U test.

Taken together, our results indicate that PNFlx treatment influences the formation of circuit stabilizing PNNs in the hippocampus during critical period windows wherein the neurocircuitry undergoes substantial experience-dependent fine tuning. The changes arise in the absence of any major effect on the total numbers of hippocampal PV-positive, CR-positive, SST-positive and Reelin-positive interneurons, with the exception of a small but significant increase in SST neuron number in the DG subfield of PNFlx animals in adulthood.

### 1.3.3 *Adult animals with a history of postnatal fluoxetine treatment exhibit a decline in hippocampal GABA-A*a*2 receptor subunit expression and enhanced c-Fos expression*

PNNs are known to prevent the lateral movement and exchange of transmembrane receptor molecules (Sorg et al., 2016). Given we noted changes in PNN numbers following PNFlx treatment accompanied by gene expression changes in ECM-associated molecules, we sought to address whether receptor-subunit switches of the NMDA and GABA receptors, associated with maturation of the hippocampal circuitry (Davis et al., 2000; Gambrill & Barria, 2011), are disrupted as a result of PNFlx treatment. We examined two classic maturation associated receptor-subunit switches, namely the levels of hippocampal expression of NR2B and NR2A, and GABA-Aα1 and GABA-Aα2, in vehicle- and PNFlx-treated adult animals. Western blotting for the proteins of interest revealed that while NR2A, NR2B and GABA-Aa1 expression levels in the hippocampus remain unaffected, GABA-Aa2 expression was significantly reduced within the hippocampi of adult animals with a history of PNFlx treatment (Fig 5B-D) (*p* value = 0.033, n = 4 per group).

**Figure 5.**
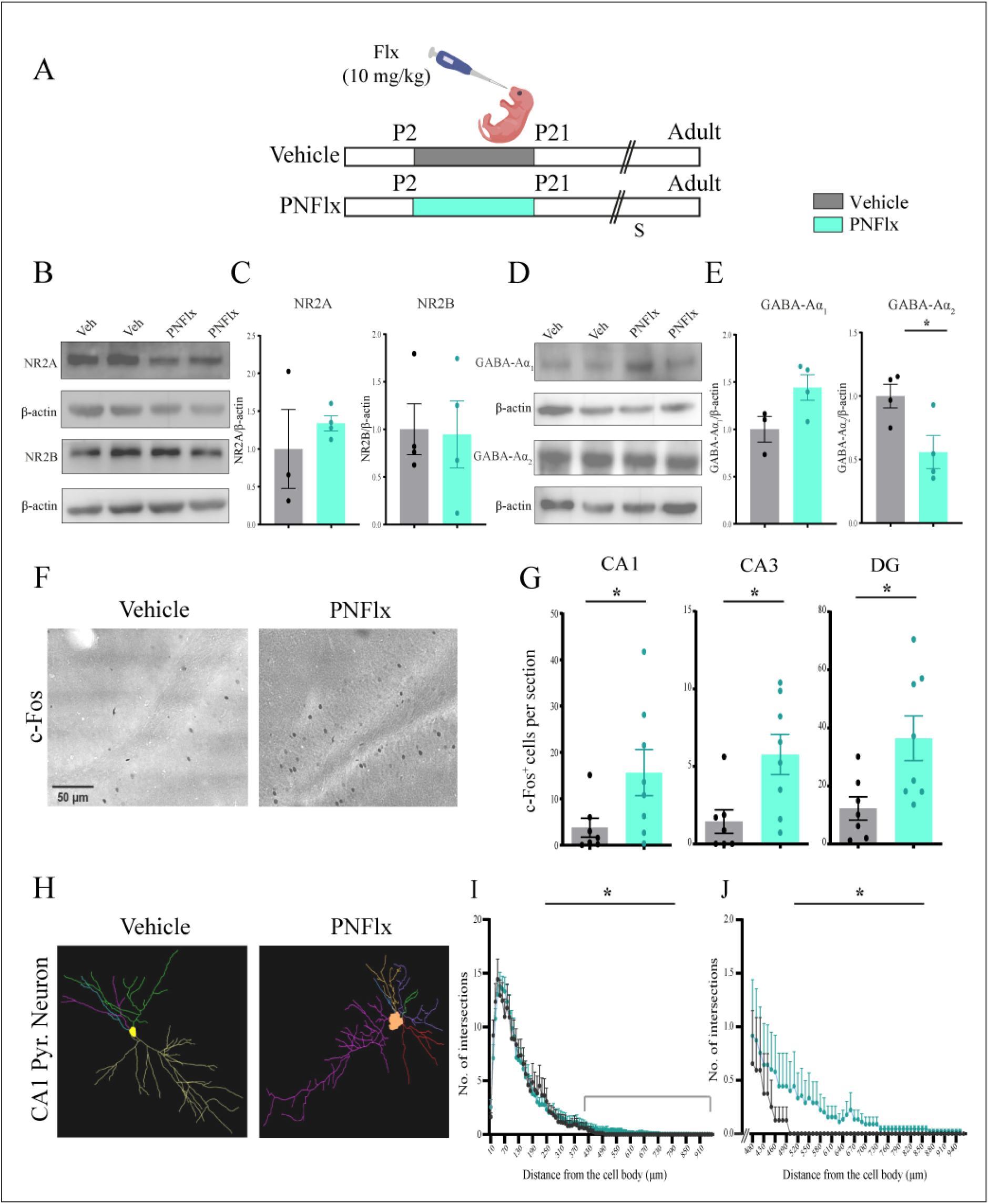
Influence of postnatal fluoxetine treatment on NMDA and GABA receptor subunit expression, c-Fos-positive cell numbers and CA1 pyramidal neuron morphology in the adult hippocampus. (A) Shown is a schematic depicting the experimental paradigm to test the effects of PNFlx administration from P2 to P21 on hippocampal NMDA and GABA receptor subunit expression, c-Fos expression, and architecture of CA1 pyramidal neurons in adult animals. S denotes the age (P80-100) at which animals were sacrificed. (B) Shown are representative western blots of NR2A and NR2B with ß-Actin as loading control from the hippocampus of vehicle-treated and PNFlx-treated adult animals. (C) Shown are densitometric quantifications of normalized protein levels of NR2A and NR2B. Normalized levels of NR2A and NR2B showed no difference between vehicle- and PNFlx-treated animals in adulthood (D) Shown are representative western blots of GABA-Aα1 and GABA-Aα2 with ß-Actin as loading control from the hippocampus of vehicle- and PNFlx-treated adult animals. (E) Shown are densitometric quantification of normalized protein levels of GABA-Aα1 and GABA-Aα2. Normalized levels of GABA-Aα2 showed a significant increase in the hippocampus of adult PNFlx-treated adult animals compared to vehicle treated adult animals while no significant difference was seen in GABA-Aα1 between vehicle and PNFlx-treated animals in adulthood. (F) Shown are representative images of c-Fos expressing cells in the hippocampus from vehicle treated and PNFlx-treated animals in adulthood. (G) Quantifications of c-Fos expressing cells showed a significant increase within the hippocampal subfields of CA1, CA3 and DG in adult PNFlx-treated animals. (H) Shown are representative traces of Golgi-Cox stained CA1 hippocampal pyramidal neurons of vehicle-treated and PNFlx-treated adult animals. (I) Shown is quantification of the number of intersections per μm distance from the soma across the entire apical dendritic arbor in Golgi-Cox stained hippocampal CA1 pyramidal neurons of PNFlx- and vehicle-treated adult animals. (I) Shown is the quantification of total number of intersections in Golgi-Cox stained hippocampal CA1 pyramidal neurons of PNFlx- and vehicle-treated adult animals. (J) Shown is quantification of the number of intersections per um distance from the soma across the distal end (400-940 μm) of the apical arbors of Golgi-Cox stained hippocampal CA1 pyramidal neurons of PNFlx- and vehicle-treated adult animals. Results for western blots, c-Fos immunohistochemistry, total number of intersections of Golgi-Cox stained CA1 pyramidal neurons are expressed as the mean ± SEM, \**p*<0.05 as compared to vehicle treated rats using two-tailed, unpaired Student’s *t-*test. Results for number of intersections and number of intersections per um distance from the soma of Golgi-Cox stained CA1 pyramidal neurons are expressed as the mean ± SEM and PNFlx animals have been compared to vehicle treated animals using non-parametric repeated measures Friedman test.

We then addressed whether the altered PNN formation, perturbed gene expression of ECM-associated molecules and dysregulation of GABA-Aα2 expression in the hippocampi of PNFlx animals was also associated with any change in baseline neuronal activity levels by assessing the influence of PNFlx on expression of the neuronal activity marker, c-Fos (Bullitt, 1990). Analysis of c-Fos expression in the hippocampus revealed that adult animals with a history of PNFlx treatment showed significantly higher numbers of c-Fos-positive cells in the hippocampal CA1 (*p* value = 0.018, n = 7 per group), CA3 (*p* value = 0.011, n = 7 per group) and DG (*p* value = 0.017, n = 7 per group) hippocampal subfields (Fig 5F, G). These observations indicate that baseline neuronal activity within the hippocampus is increased in adult animals with a history of PNFlx treatment.

Given that neuronal architecture is known to be strongly correlated with changes in neuronal activity, and is also dependent on the surrounding ECM (Levy et al., 2014), we addressed whether PNFlx treatment led to persistent changes in dendritic morphology of the principal CA1 pyramidal neurons. To address this, we utilized Golgi-Cox staining to trace the processes of CA1 pyramidal neurons in vehicle- and PNFlx-treated adult animals. Sholl analysis revealed a small, but significant, increase in dendritic arbors at the distal-most ends of the apical branches of CA1 pyramidal neurons as a consequence of PNFlx treatment (Fig 5H-J) (*p* value < 0.001, n = 8 - 9 per group). We have not examined the influence of PNFlx exposure on spine density, or differences in morphology that may be noted at earlier time points.

Collectively, our results indicate that PNFlx treatment evokes a delayed development of PNNs in the hippocampus during the postnatal window and results in persistent changes in neuronal activity, GABA receptor subunit composition and CA1 pyramidal neuron dendritic complexity that are noted in adulthood long after the cessation of fluoxetine treatment.

## Discussion

Here, we have performed a detailed characterization of the short- and long-term consequences of PNFlx exposure on PNNs, hippocampal interneuron number and regulation of ECM-related gene expression. We find a significant reduction in PNN numbers within the CA1 and CA3 hippocampal subfields, immediately following the cessation of PNFlx treatment at P21, and this decline persists within the CA1 subfield into adulthood. We did not observe any change in the numbers of specific interneuron classes, characterized by the expression of PV, SST, CalR and Reelin, in the hippocampus at P21. However, we did note a small, but significant, increase in SST-positive interneurons in the DG hippocampal subfield in adulthood in the PNFlx cohort. Our results suggest that PNFlx disrupts the normal ontogeny of the hippocampus, which is known to exhibit a protracted window of development extending well into postnatal life and adolescence (Bayer, 1980; Rice & Barone, 2000). This indicates that serotonin elevation via PNFlx may result in a delayed closure of critical period plasticity in the hippocampus, revealed via the delay in the development of PNNs, in particular those that ensheath PV-positive interneurons. Given that the temporal window of PNFlx overlaps with substantial neuronal plasticity in terms of interneuron migration, apoptosis, synaptic pruning and synaptogenesis, as well as the formation of PNNs, our results provide an insight into the developmental consequences of enhanced serotonin levels on the maturing hippocampal neurocircuit (Gingrich et al., 2017). Further, we show that PNFlx treatment leads to a reduction in hippocampal GABA-Aa2 receptor subunit protein levels, and an increase in c-Fos expression across all hippocampal subfields in adulthood, suggesting persistent dysregulation of neuronal activity in the hippocampus as a consequence of the early life fluoxetine exposure. This is accompanied by increased dendritic complexity in the distal regions of the apical branches of CA1 pyramidal neurons in adult animals with a history of PNFlx. Collectively, our findings reveal both early-onset and persistent changes in PNN formation, interneuron number, neuronal activity, CA1 pyramidal neuron cytoarchitecture and altered transcriptional regulation of ECM and PNN-associated genes in the hippocampus following PNFlx treatment (Fig 6).

**Figure 6.**
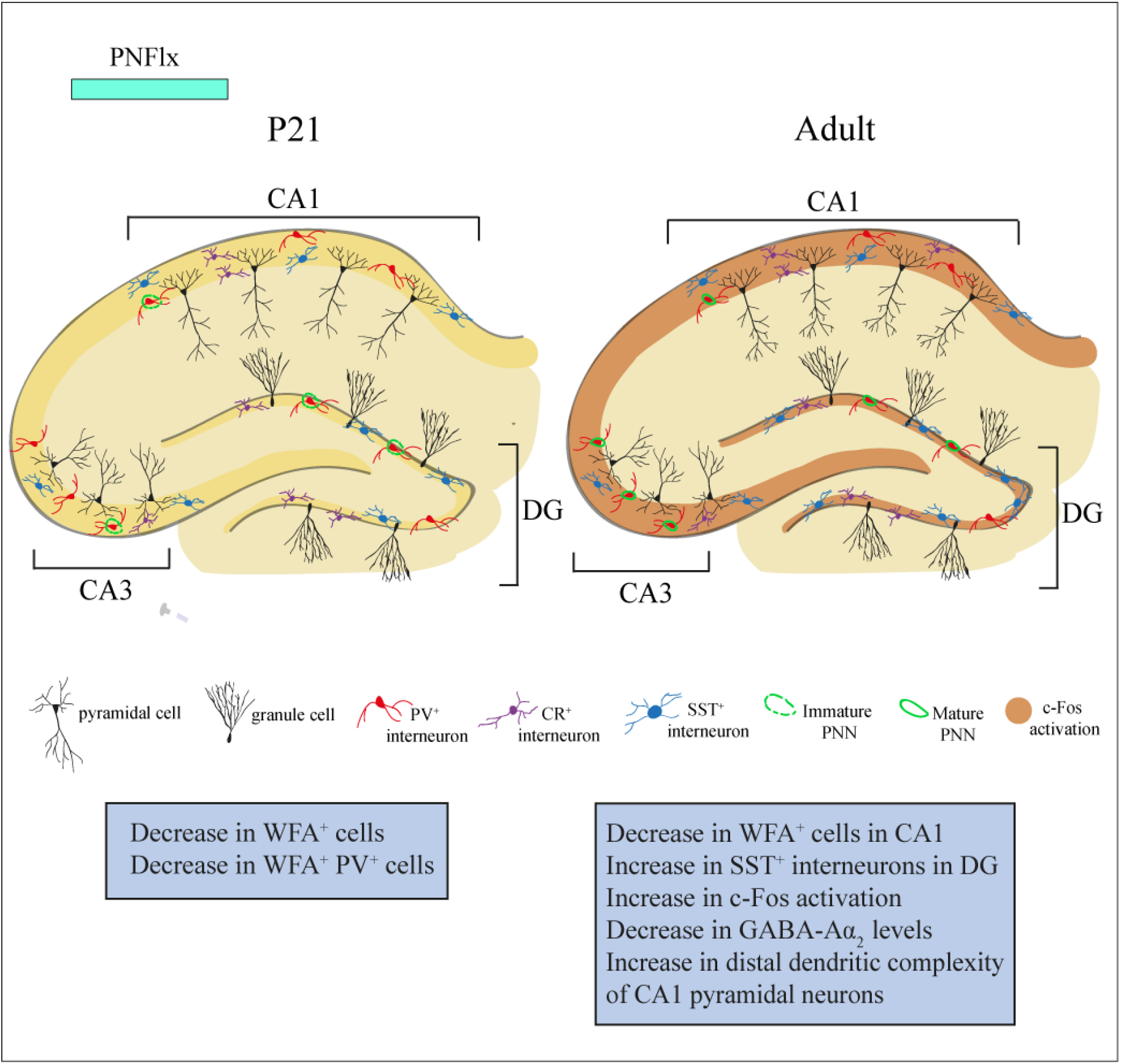
Postnatal fluoxetine treatment causes immediate and long-lasting cellular and molecular changes in the rat hippocampus. PNFlx treatment alters the developmental trajectory of PNN formation within the hippocampus. PNFlx treatment evoked a decline in the number of WFA-positive PNNs, and PV-positive cells ensheathed by WFA-positive PNNs, in the CA1 and CA3 hippocampal subfields at P21. A history of PNFlx treatment was associated with a decline in the number of WFA-positive PNNs in the CA1 hippocampal subfield, and a small increase in the number of SST-positive neurons in the DG hippocampal subfield in adulthood. Adult animals with a history of PNFlx also exhibited an increase in the number of c-Fos positive cells in all hippocampal subfields, a decline in GABA-Aα2 receptor subunit expression within the hippocampus, and enhanced dendritic complexity noted in the distal dendrites of the apical branches of CA1 pyramidal neurons.

Serotonin modulates the development and maturation of neurons during early life (Gaspar et al., 2003). Perturbations in serotonin levels or signaling during this period can lead to developmental, cytoarchitectural as well as behavioral changes that last well into adulthood (Ansorge et al., 2004; Brummelte et al., 2017; Shah et al., 2018). Early-life SSRI administration has been suggested to prolong the plasticity period as examined via delayed PNN formation in multiple limbic circuits (Kobayashi et al., 2010; Umemori et al., 2015). A previous study showed that administration of fluoxetine in drinking water to pregnant mouse dams, commencing gestational day 7 and extending until P7, which would presumably result in fluoxetine exposure to nursing pups, showed a significant reduction in PNN numbers in the CA1 hippocampal subfield at P17, and exhibited recovery in cohorts of pups examined at P24 (Umemori et al., 2015). While the present study differs substantially in the mode of fluoxetine administration, treatment timeline, species studied and ages examined, it is note-worthy that we also find a robust decline in PNN numbers at the early time-point of P21 following PNFlx treatment. At the P21 timepoint following PNFlx we have examined PNN numbers, as well as the number of PV-positive interneurons decorated by PNNs, in both male and female pups. However, our studies in adulthood are restricted to males with a history of PNFlx administration, and this is one of the caveats of our study which does not allow us to further address potential sexual dimorphism in the effects of a history of PNFlx in adulthood. Studies have also shown that chronic administration of SSRIs in adulthood evokes a juvenile-like plasticity in the visual and prefrontal cortex associated with a dissolution of PNNs (Guirado et al., 2014; Ohira et al., 2013, 2019; Vetencourt et al., 2008). PNN formation, primarily around PV-positive interneurons, is thought to be a critical stage that defines the establishment of the maturity, and closure of critical-period plasticity, which emerges at distinct developmental epochs in a circuit-specific manner (Köppe et al., 1997; Lensjø et al., 2016). Our findings indicate that the delayed appearance of PNNs following PNFlx treatment results in significantly fewer numbers of PV-positive interneurons ensheathed by PNNs in the CA1 and CA3 hippocampal subfields at P21. This decline is transient, and by adulthood the number of PV-positive interneurons decorated with PNNs is indistinguishable between vehicle- and PNFlx-treated animals. However, our results do not allow us to resolve whether the constituent components of the PNN at these timepoints are perturbed. PNNs play an important role in stabilization of synaptic contacts, structural and functional plasticity, and the establishment of excitation/inhibition (E/I) balance in neural circuits (Fawcett et al., 2019; Hensch, 2005). Our results clearly indicate that the establishment of PNNs in specific hippocampal subfields is disrupted as a consequence of PNFlx treatment, and that this disruption of PNNs persists into adulthood in a hippocampal subfield-specific manner.

Gene expression analysis revealed that the hippocampal expression of several PNN and ECM-associated genes such as *aggrecan*, *versican*, *brevican*, *neurocan*, *cspg4* and *hapln1-2* do not exhibit any change in expression at either P21 or adulthood following PNFlx treatment. We did however, observe a decrease in *mmp2* transcript levels immediately post-cessation of the PNFlx treatment at P21, and also noted a long-term decline in *tenascin c* and *has3* mRNA within the hippocampus in adulthood in animals with a history of PNFlx administration. While we cannot draw direct causal links between the transcriptional dysregulation of individual genes to the complex cellular and behavioral phenotypes noted with PNFlx, it is interesting to note that mice constitutively deficient in tenascin C display compromised LTP in CA1 pyramidal neurons (Evers et al., 2002), and mutant mice that were combinatorial knock-outs of *tenascin C*, t*enascin R*, *brevican* and *neurocan* were shown to have decreased PNN area and intensity in the CA2 hippocampal subfield (Gottschling et al., 2019). Analyses of interneuron numbers revealed that while the numbers of PV-, CalR-, and Reelin-positive neurons in the hippocampus do not change as a consequence of PNFlx treatment, we do find a significant increase in the number of SST-positive neurons in the DG of adult animals with a history of PNFlx. Multiple studies indicate that perturbed SST-signaling, and a loss of SST-positive neurons are both linked to elevated depressive-/anxiety-like behaviors, and vulnerability of SST neurons to damage has also been suggested in the context of major depressive disorder (Douillard-Guilloux et al., 2017; Fee et al., 2017; Lin & Sibille, 2015). Hippocampal SST neurons are also linked to the control of HPA axis activity, with differential effects observed on emotionality based on the class of SST receptors involved (Prévôt et al., 2017; Scheich et al., 2016; Yamamoto et al., 2018). Given the complexity of information processing in hippocampal microcircuitry, it is unclear at present what the consequence of a small increase in SST-neuron number in the DG would be on hippocampal function. PNFlx treatment does not appear to result in major disruption of interneuron numbers during postnatal life or in adulthood, however we cannot preclude effects on the cytoarchitecture and function of specific interneuron classes. The decline in PNNs, in particular those that encapsulate PV-positive interneurons, following PNFlx treatment raises the possibility of a disruption of the intrinsic properties of PV-positive interneurons, and hence an impact on the ability of PV-positive interneurons to dynamically gate and adapt their responses to altered neuronal activity (Favuzzi et al., 2017).

Disruption of PNN formation and ECM abnormalities have been linked to the pathophysiology of several psychiatric disorders, including schizophrenia and mood disorders and neurodevelopmental disorders, such as autism spectrum disorder and epilepsy (Pantazopoulos & Berretta, 2016; Wen et al., 2018). Preclinical models of early life trauma that are based on the disruption of dam-pup interactions, such as maternal separation (MS) and limited nesting bedding (LNB), perturbed inflammatory milieu evoked via maternal immune activation (MIA) and exposure to juvenile stress have all been reported to evoke a disruption of PNN formation. MS is reported to result in a reduced density of PNNs in the prelimbic cortex at P20 (Gildawie et al., 2020), and male pups raised by dams in the LNB paradigm exhibit a sexually dimorphic increase in PNN density around PV-positive interneurons within the right basolateral amygdala (Guadagno et al., 2020). Pups born to dams subjected to MIA showed a decline in the density of PNNs in the basolateral amygdala at P35, and in the prelimbic cortex in adulthood (Paylor et al., 2016). Exposure to juvenile stress is linked to a decrease in the intensity of PNN staining in the CA1 hippocampal subfield, the dorsal anterior cingulate cortex, the infralimbic cortex and the motor cortex, immediately following the cessation of the stress (Ueno et al., 2018). Interestingly, MS, LNB, MIA and juvenile stress are all reported to evoke a perturbation of serotonergic signaling (Benekareddy et al., 2010; Goeden et al., 2016; Jablonski et al., 2017; Luo et al., 2015). These results raise the intriguing possibility that both a perturbation of serotonergic signaling, as well as an altered trajectory of PNN development in key limbic circuits, may be associated with the development of anxiogenic and depressive-like behavioral phenotypes noted in adulthood in these models of early adversity. Our studies suggest that PNFlx treatment, which evokes persistent increases in anxiety- and despair-like behavior across the life-span, also targets the formation of PNNs in the hippocampus, a brain region implicated in the regulation of mood-related behavior. A disruption of molecular and cellular regulators of critical period plasticity during postnatal maturation has been suggested to be a common signature associated with animal models of depression, based on early adversity exposure, as well as genetic and pharmacological models of neurodevelopmental disorders (Callaghan et al., 2013; Do et al., 2015; Fagiolini & Leblanc, 2011; Spijker et al., 2020).

PNN formation and maturation marks the end of the critical period plasticity window in most circuits, and a delay in PNN maturation is suggestive of a protracted closure of the critical period (Hensch, 2005). A shift in the receptor subunit composition of both NMDA and GABAA receptor subunits is a hallmark signature associated with neurodevelopment and with the establishment of E/I balance (Davis et al., 2000; Gambrill & Barria, 2011). We find that while the ratio of NR2A/NR2B receptor subunits is not altered in adult animals with a PNFlx history, we note a robust decline in GABA-Aα2 receptor subunit in the hippocampi of PNFlx-treated animals in adulthood. The natural developmental progression indicates a switch from GABA-Aα2 to GABA-Aα1 receptor subunits in adults (Fritschy et al., 1994), and a change in the ratio of specific GABA-A receptor subunits following PNFlx treatment could have an important implication for neuronal activity within maturing hippocampal circuitry. Indeed, this is reflected in the c-Fos expression analysis which indicates an increase in expression of the neuronal activity marker in all hippocampal subfields of PNFlx animals in adulthood. Perturbed baseline expression of an immediate early gene marker noted in adulthood in animals with a history of PNFlx administration points to potential disruption of neuronal activity. Our findings motivate further studies to directly test the consequences of PNFlx-evoked altered PNN development on the neuronal activity and intrinsic properties of PV-positive interneurons, and on the emergence of E/I balance within the hippocampus.

Serotonin is known to modulate neuronal morphology, and dendritic spine shape/density (Bijata et al., 2017; Daubert & Condron, 2010; Wirth et al., 2017). PNFlx treatment from P2 to P11 results in decreased dendritic arbor complexity in pyramidal neurons of the infralimbic cortex (Rebello et al., 2014). In contrast, a developmental knockout of the serotonin transporter is associated with increased dendritic complexity in infralimbic pyramidal neurons, and an increase in spine density in pyramidal neurons of the basolateral amygdala (Wellman et al., 2007). We noted a subtle increase in dendritic complexity, restricted to the distal-most regions of the apical dendrites of CA1 pyramidal neurons in PNFlx-treated animals in adulthood. Hippocampal gene expression changes associated with PNFlx indicate dysregulation of pathways that modulate neuronal cytoarchitecture, namely mTOR signaling, and perturbed gene expression of the hyperpolarization-activated cyclic nucleotide-gated 1 channel (*Hcn1*) which is known to exhibit distal dendrite enrichment (Sarkar et al., 2014). While we have not extensively examined the influence of PNFlx on spine number, shape and density in hippocampal pyramidal neurons, our observations motivate a detailed characterization of the impact of early life serotonin elevation on the cytoarchitecture of distinct neuronal classes within the hippocampus.

Collectively, our findings indicate that early life elevation of serotonin via treatment with the SSRI fluoxetine alters the developmental trajectory of PNNs within the hippocampus, suggestive of a disruption of critical period plasticity. This evidence supports an emerging body of literature that implicates disruption of PNNs and ECM dysregulation as putative mechanisms that contribute to the circuit dysfunction associated with psychiatric and neurodevelopmental disorders.

## Acknowledgements

We thank Dr. Shital Suryavanshi, Ms. K.V. Boby and the animal house staff at the Tata Institute of Fundamental Research (TIFR), Mumbai for technical assistance.

